# Influenza A virus coinfection dynamics are shaped by distinct virus-virus interactions within and between cells

**DOI:** 10.1101/2022.11.27.515562

**Authors:** Gabrielle K. Delima, Ketaki Ganti, Katie E. Holmes, Jessica R. Shartouny, Anice C. Lowen

## Abstract

When multiple viral populations propagate within the same host environment, they often shape each other’s dynamics. These interactions can be positive or negative and can occur at multiple scales, from coinfection of a cell to co-circulation at a global population level. For influenza A viruses (IAVs), the delivery of multiple viral genomes to a cell substantially increases burst size. However, despite its relevance for IAV evolution through reassortment, the implications of this positive density dependence for coinfection between distinct IAVs has not been explored. Furthermore, the extent to which these interactions within the cell shape viral dynamics at the level of the host remains unclear. Here we show that, within cells, coinfecting IAVs strongly augment the replication of a focal strain, irrespective of the native host of the coinfecting IAV or its homology to the focal strain. Coinfecting viruses with a low intrinsic reliance on multiple infection offer the greatest benefit. Nevertheless, virus-virus interactions at the level of the whole host are antagonistic. This antagonism is recapitulated in cell culture when the coinfecting virus is introduced several hours prior to the focal strain or under conditions conducive to multiple rounds of viral replication. Together, these data suggest that beneficial virus-virus interactions within cells are counterbalanced by competition for susceptible cells during viral propagation through a tissue. The integration of virus-virus interactions across scales is critical in defining the outcomes of viral coinfection.

## Introduction

The past three influenza A virus (IAV) pandemics arose through reassortment involving human and non-human IAVs in coinfected hosts (1–6). Due to the segmented nature of the IAV genome, when two or more IAVs coinfect the same cell, viral progeny can contain a mix gene segments from either parental strain (7–9). Reassortment between phylogenetically distant IAVs often generates progeny less fit than the parental strains due to the unlinking of coevolved gene segments (10, 11). However, reassortment can facilitate host switching through the formation of an antigenically novel strains that carry gene segments well-adapted to human infection(2, 12). Thus, coinfections involving heterologous IAVs are of constant concern.

When viruses coinfect the same host or cell, each can affect the other’s replication. These virus-virus interactions range from antagonistic to beneficial, depending on the context. At the cellular scale, viruses may compete directly for limited resources within the cell or indirectly through the triggering of antiviral responses. These antagonistic effects are typically potent in the case of defective viral genomes(13). Conversely, coinfection may enhance productivity through complementation of incomplete or otherwise defective viral genomes or by increasing the availability of viral proteins needed for replication or host immune suppression(14–17). At the host scale, viruses may compete for naïve cells: in a mechanism known as superinfection exclusion, infected cells can become refractory to secondary infection. Additionally, activation of non-specific antiviral responses can further suppress viral infection within the host(18). Conversely, a focal virus may benefit indirectly from the suppression of systemic antiviral responses or induction of pro-viral processes (e.g. coughing) by a coinfecting virus. These virus-virus interactions often occur simultaneously, but which interaction is dominant may vary with context.

Our prior work focused on virus-virus interactions between homologous viruses. We previously reported that coinfection of a cell with multiple IAV particles is often required to initiate a productive infection(15). Furthermore, we determined that this reliance on multiple infection is a common feature of IAVs, but the level of reliance is both strain and host dependent(16). Density dependence of IAV replication arises due to a need for complementation of incomplete viral genomes (15) and can be heightened in the context of deleterious mutation or antiviral drug treatment (17). It follows that interactions between coinfecting IAVs impact the likelihood of productive vs abortive infection. Thus, IAV virus-virus interactions are a major factor defining the outcomes of cellular infection. These outcomes include whether or not progeny are produced, their quantitative yield and their genotypes. Given that the replicative potential of a virus is strongly shaped by its interactions with other co-occurring viruses, we sought to understand the implications of virus-virus interactions in the context of coinfection between phylogenetically distinct IAVs.

Here, we investigate what viral traits define the extent to which a coinfecting IAV can augment the replication of another. Specifically, we examine the importance of a coinfecting virus’ intrinsic reliance on multiple infection, the host species to which it is adapted, and its homology to the focal virus. To this end, we evaluated the outcomes of coinfection with a set of well-characterized IAV strains: influenza A/guinea fowl/Hong Kong/GFHK99/99 (H9N2) (GFHK99), A/mallard/Minnesota/199106/99 (H3N8) (MaMN99), A/Netherlands/602/2009 (H1N1) (NL09), and A/Panama/2007/99 (H3N2) (Pan99) viruses. Our results indicate that homology and host adaptation are not critical for defining IAV-IAV interactions within cells. Conversely, IAVs having a low intrinsic reliance on coinfection more effectively augment replication of the focal virus. While these beneficial interactions are readily detected when infection is limited to a single viral generation, under conditions that allow further propagation, antagonistic effects become predominant. Dynamics in vivo and during multiple rounds of viral infection in cell culture suggest that competition for target cells and super infection exclusion typically limit the opportunity for distinct viral populations to interact within cells.

## Results

### Intrinsic reliance on multiple infection in mammalian cells is strain dependent

Previously we showed that influenza A virus replication is strongly dependent on viral density. Namely, progeny production is enhanced when population density is sufficient to ensure delivery of many viral genomes to a cell. The extent of this dependence was, however, noted to vary with virus strain. To investigate the importance of this phenotype for interactions between distinct coinfecting strains, we sought to identify a panel of viruses that vary in their reliance on multiple infection. To this end, we evaluated the sensitivity of viral RNA (vRNA) replication to multiplicity of infection (MOI) for a diverse set of IAV strains, including two of avian-origin and two of human-origin. This was done by infecting Madin Darby canine kidney (MDCK) cells with a wild type (wt) virus at a constant low dose and infecting simultaneously with increasing doses of a homologous virus, termed var. The var virus carries synonymous mutations to allow differentiation of its genome from that of the corresponding wt virus. Coinfections were limited to a single cycle of infection to ensure that results reflected processes occurring within cells. In line with previous observations, GFHK99wt virus replication is enhanced ~100-fold by co-inoculation with GFHK99var virus (Fig 1A). However, the replication of a second avian virus, MaMN99wt shows at most a 4-fold enhancement with addition of MaMN99var (Fig 1B). While peak enhancement of Pan99wt virus with the addition of Pan99var virus is ~900-fold (Fig 1C), NL09wt replication is reduced approximately 300-fold through the addition of NL09var (Fig 1D). These data indicate that reliance on multiple infection is strain dependent and does not consistently correspond to the extent of viral adaptation to the host.

**Fig 1.**
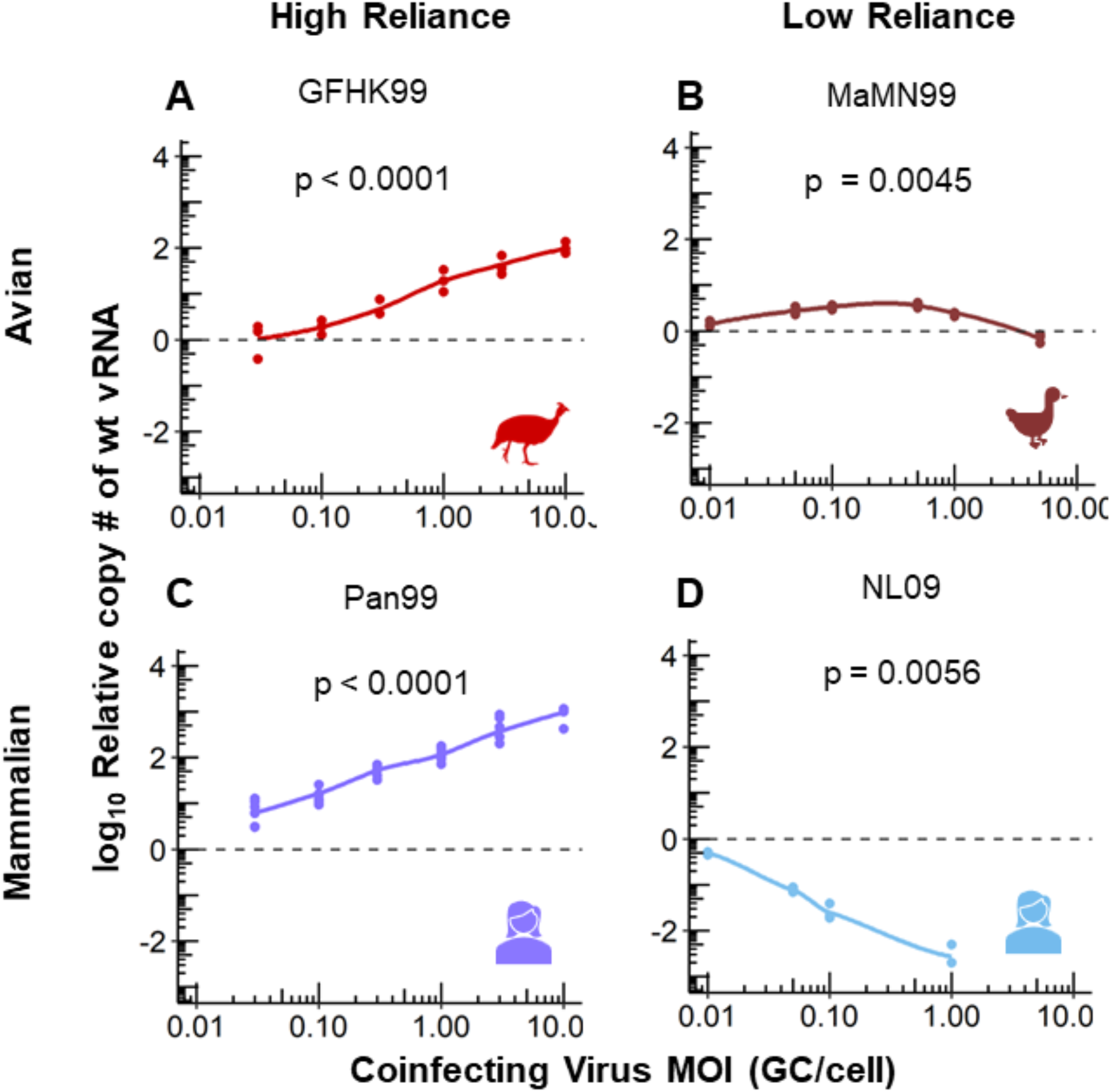
Intrinsic reliance on multiple infection in mammalian cells is strain dependent. MDCK cells were coinfected with homologous wt/var viruses of the indicated strain backgrounds. The wt viruses had an MOI of 0.005 genome copies (GC)/cell and coinfecting var viruses were added with increasing MOI ranging from 0.005 to 10 GC/cell. The fold change in wt vRNA copy number, relative to wt only infection (dashed line), is plotted. Results of three to six biological replicates derived from one to two independent experiments are plotted and solid lines connect the means of the points shown. Each line was evaluated by linear regression to determine if the slope was significantly non-zero. Data in panels B and D were reported in (16).

### Beneficial interactions within cells extend to heterologous virus pairings

To date, our examination of IAV density dependence has centered on homologous interactions that would occur within a given viral population. Here, we sought to test whether the positive effects of increased density would extend to coinfection with distinct IAV strains. Although virus-virus interactions would typically be bi-directional, for the purposes of our experimental design, we measure the impact of coinfection on only one strain in the pairing, referred to here as the focal virus.

We predicted that specific traits of a coinfecting virus, such as the strength of its intrinsic density dependence, the host to which it is adapted, or its degree of homology to the focal strain, may affect its potential to modulate the replication of the focal strain. To test these predictions, we performed a series of coinfections using GFHK99wt as the focal virus and MaMN99, NL09, or Pan99 as coinfecting viruses. We infected MDCK cells with a constant dose of GFHK99wt virus and increasing doses of GFHK99var virus (as a control) or NL09, MaMN99, or Pan99 virus, then quantified levels of GFHK99wt genomes in the cells. These coinfections were again limited to a single cycle to ensure that results reflected processes occurring within cells. The results show that GFHK99wt genome levels increase with increasing doses of every coinfecting var virus, indicating that phylogenetic relatedness within the species *influenzavirus A* is not required for beneficial interactions within the cell (Fig 2 and S1 Fig). However, the degree to which GFHK99wt virus replication was increased varied with the coinfecting strain. Compared to the control in which GFHK99var was the coinfecting virus, MaMN99 and NL09var virus coinfections resulted in significantly greater enhancement of GFHK99wt virus replication (Fig 2A,C and S1 Fig) while coinfection with Pan99var virus resulted in lower enhancement (Fig 2B). These results indicate that the degree to which a coinfecting virus is adapted to the host (in this case, mammalian cells) is not a major factor defining its potential to augment replication of the focal virus. Instead, the data suggest that lower intrinsic density dependence of the coinfecting strain, as for NL09 and MaMN99, allows a stronger benefit to be conferred on the focal virus.

**Fig 2.**
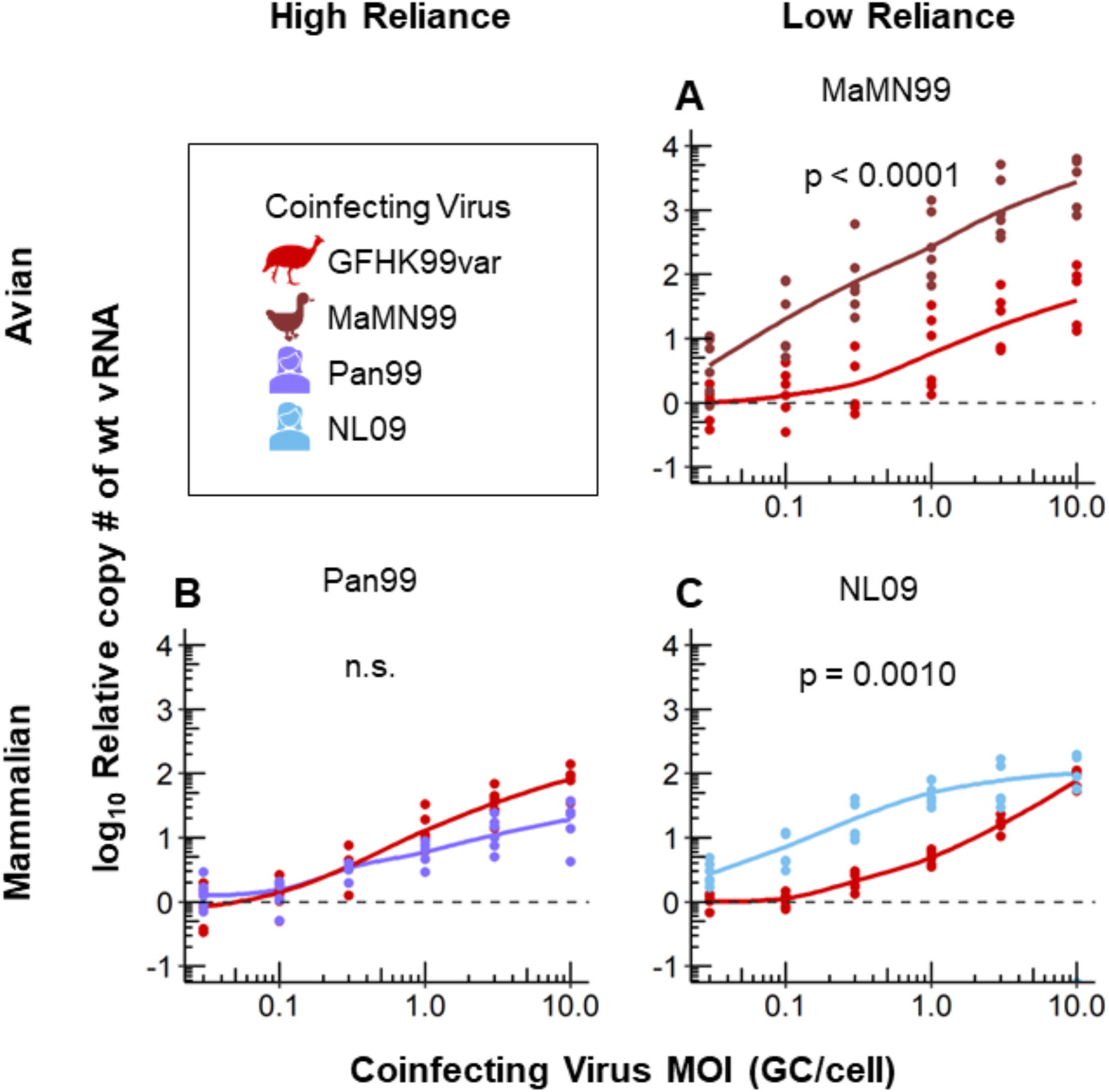
Coinfection with a virus with low reliance on multiple infection results in a greater increase in GFHK99 virus replication. MDCK cells were coinfected with GFHK99wt virus at an MOI of 0.005 GC/cell and increasing doses of the homologous GFHK99var virus (red in each panel) or a heterologous MaMN99 (A), NL09 (B), or Pan99 (C) virus. The fold change in GFHK99wt vRNA copy number, relative to GFHK99wt-only (dashed line), is plotted. Results of six biological replicates derived from two independent experiments are plotted and solid lines connect the means. Significance of differences between coinfecting viruses was evaluated by two-way ANOVA.

### Coinfection with a virus with low reliance on multiple infection better meets IAV need for help

To more rigorously examine how the intrinsic reliance on multiple infection displayed by a coinfecting virus modulates its impact on a focal virus, we used a mutant strain of GFHK99 virus. We previously showed that introduction of a PA K26E mutation into the GFHK99 strain reduces reliance on multiple infection(17). We confirmed this phenotype here using homologous wt and var strains of GFHK99 PA K26E (Fig 3A). We then compared the replication of GFHK99wt virus, the focal virus, during coinfection with either the homologous GFHK99var or the GFHK99 PA K26E mutant virus. We found that GFHK99wt virus genome replication was enhanced significantly more during coinfection with the low reliance GFHK99 PA K26E virus compared to coinfection with GFHK99var virus (Fig 3B). These data reinforce the role of intrinsic density dependence in defining the benefit conferred by a coinfecting strain.

**Fig 3.**
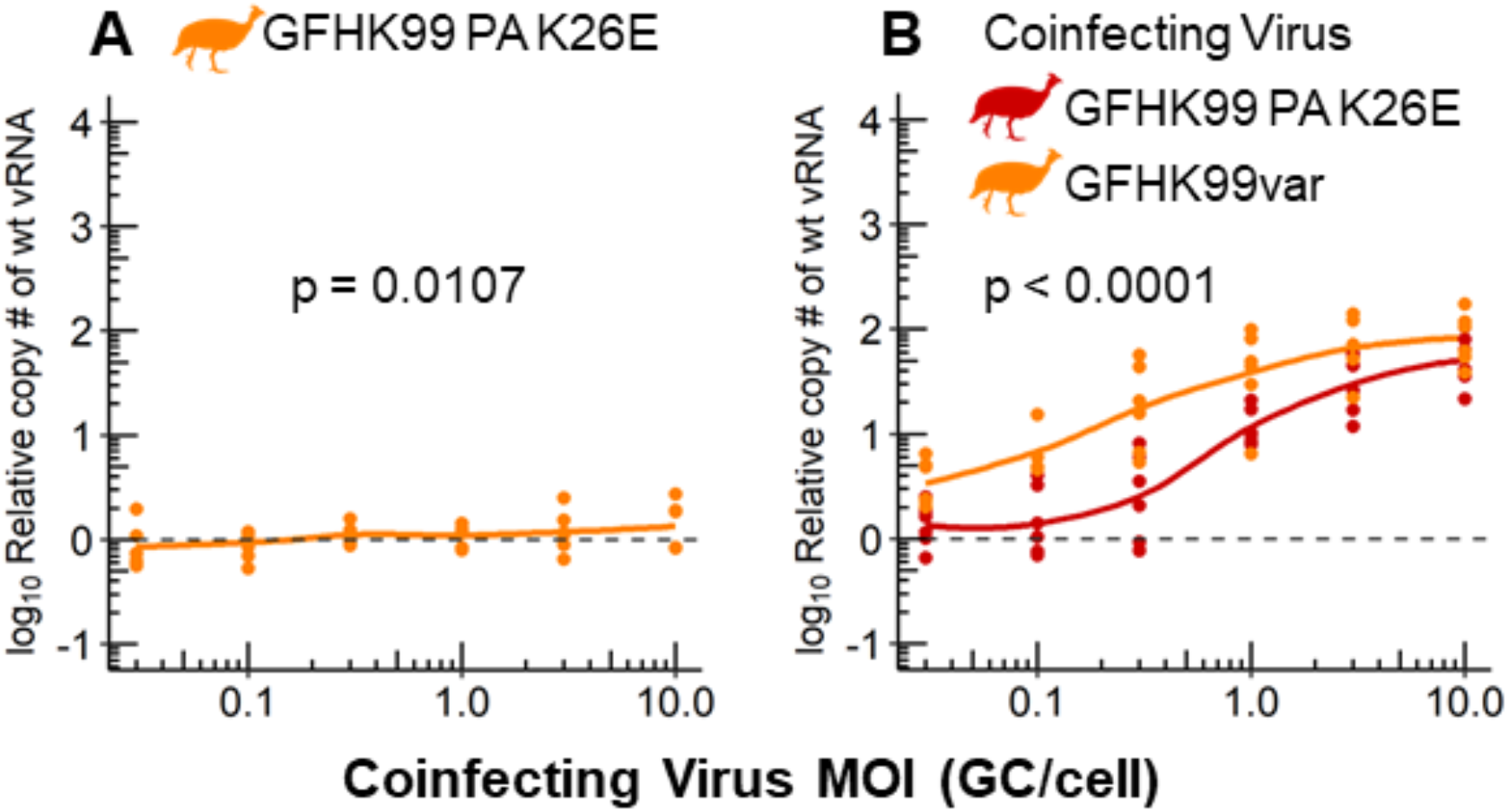
The extent to which a coinfecting virus augments replication of GFHK99 is defined by its intrinsic reliance on multiple infection. A) MDCK cells were coinfected with GFHK99wt PA K26E virus at an MOI of 0.005 GC/cell and increasing doses of a homologous GFHK99var PA:K26E virus. The fold change in GFHK99wt PA K26E vRNA copy number, relative to wt-only (dashed line), is plotted. Results of six biological replicates derived from two independent experiments are plotted and solid lines connect the means. The line was evaluated by linear regression to determine if the slope was significantly non-zero. B) MDCK cells were coinfected with GFHK99wt virus at an MOI of 0.005 GC/cell and increasing doses of GFHK99var or GFHK99var PA K26E virus. The fold change in GFHK99wt vRNA copy, relative to wt-only (dashed line), is plotted. Results of six biological replicates derived from two independent experiments are plotted and solid lines connect the means. Significance of differences between coinfecting viruses were evaluated by two-way ANOVA.

### Competitive virus-virus interactions predominate at the level of the whole host

To test the extent to which the beneficial interaction observed in our cell culture model shapes viral dynamics in vivo, we used a guinea pig model. Mimicking the experiments in cell culture, guinea pigs were inoculated simultaneously with a low dose of GFHK99wt virus and increasing doses of GFHK99var or GFHK99 PA K26E viruses. The titer of GFHK99wt virus was assessed in daily nasal wash samples. Contrary to expectation, GFHK99wt virus replication over the course of the infection was suppressed by the coinfecting virus: relative to GFHK99wt-only controls, GFHK99wt virus vRNA levels were lower during coinfection with either GFHK99var or GFHK99 PA K26E viruses (Fig 4A,B and S2 Fig).

**Fig 4.**
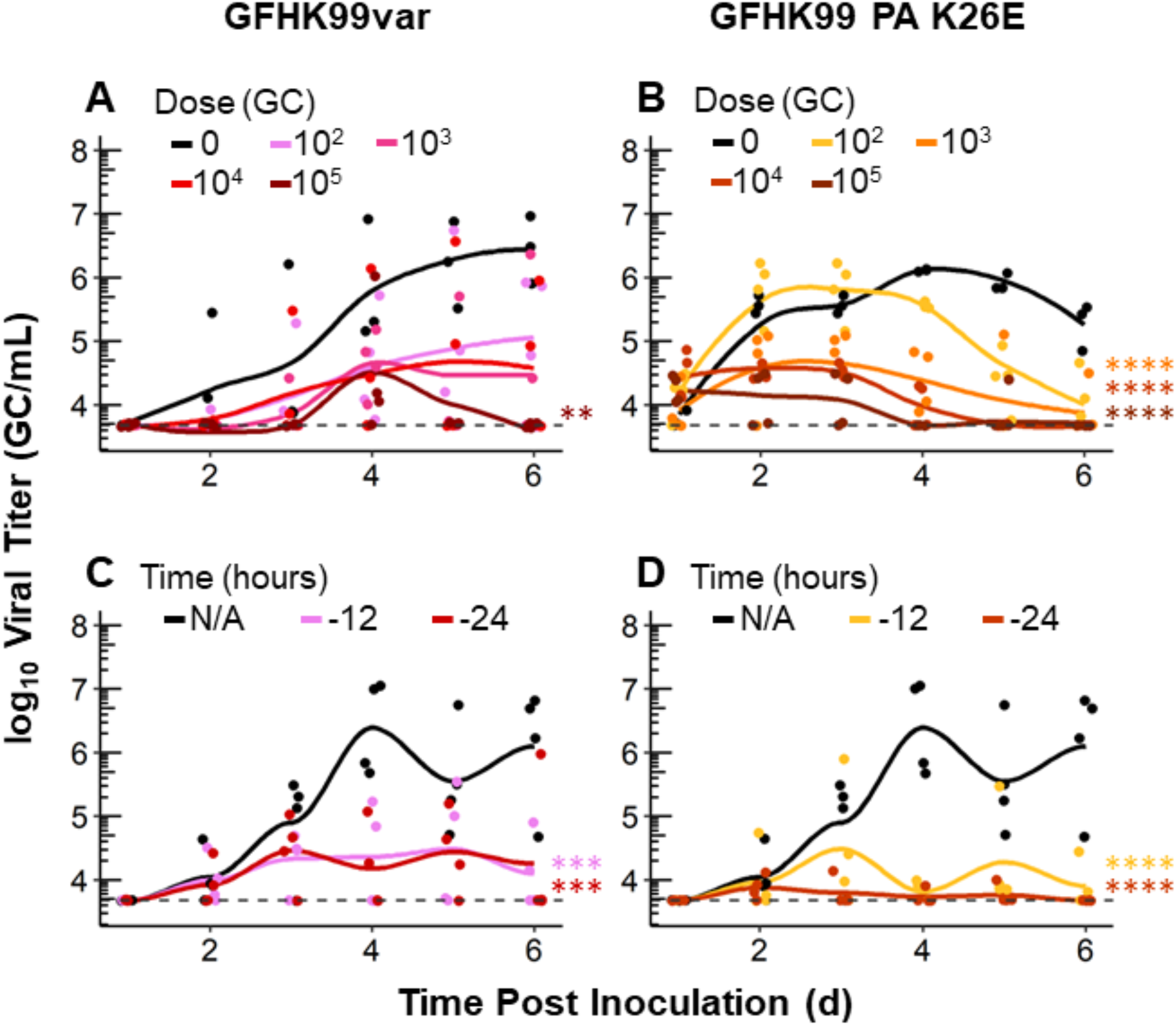
At the level of the whole host, GFHK99 replication is diminished by coinfection. A-B) Guinea pigs were coinfected with GFHK99wt at a dose of 10^3^ GC and increasing doses of the GFHK99var or GFHK99 PA K26E virus. The viral titer of GFHK99wt is plotted and the limit of is indicated by the dashed line. C-D) Guinea pigs were pre-inoculated with 10^4^ GC of either GFHK99var (C) or GFHK99 PA K26E (D) virus either 12 or 24 h prior to a dose of 10^4^ GC of GFHK99wt virus. Results of four guinea pigs per condition are plotted and solid lines connect the means. Significance of differences between coinfecting viruses were evaluated by two-way ANOVA; **p<0.01, ***p< 0.001, ****p< 0.0001.

Unlike in cell culture, multiplicity of infection cannot be controlled in vivo and mixing of coinfecting viruses can be constrained by spatial structure, limiting the potential for cellular coinfection (8, 19–21). Thus, we reasoned that the failure to detect the beneficial effects of coinfection in vivo may stem from a paucity of wt-var coinfections occurring under the conditions used. Based on prior studies (22), we hypothesized that pre-inoculation with GFHK99var or GFHK99 PA K26E viruses would increase the likelihood of cellular coinfection with the focal virus by allowing their propagation in the target tissue prior to introduction of GFHK99wt virus. Thus, in a second experiment, control guinea pigs were left alone and test guinea pigs were pre-inoculated with GFHK99var or GFHK99 PA K26E virus either 12 h or 24 h before inoculation with GFHK99wt virus, using equivalent doses. Analysis of nasal wash samples revealed that GFHK99wt virus was able to replicate in each of the four GFHK99wt-only infected guinea pigs, reaching peak titers by day 4 (Fig 4 and S3 Fig). However, GFHK99wt virus was not detected in at least one guinea pig in each group co-inoculated with GFHK99var virus and two guinea pigs in each group co-inoculated with GFHK99 PA K26E virus (Fig 4 and S3 Fig). Furthermore, in line with the previous experiment, the peak titer of GFHK99wt virus was greatest in GFHK99wt only inoculated guinea pigs (Fig 4C,D).

Taken together, these experiments revealed that the dynamics of coinfecting viruses within an intact animal host are antagonistic. Combined with our cell culture experiments, which indicate that beneficial interactions occur within individual coinfected cells, these data suggest that either the enhancement of focal vRNA replication within cells is not translated into increased viral release or that antagonistic interactions occurring at a higher spatial scale are predominant.

### Genome incorporation into progeny virions is enhanced during coinfection

To assess whether beneficial virus-virus interactions within cells results in increased virus release, we evaluated the impact of coinfection in cultured cells on the abundance of focal viral genomes in both infected cells and released progeny. Similar to the experiments outlined above, a low dose GFHK99wt virus and a high dose of a homologous var or heterologous GFHK99 PA K26E, MaMN99, or NL09 viruses were used. After 24 h, GFHK99wt virus genomes within cells and in the supernatant containing released progeny virions were quantified. Under the conditions of MOI used and at this time point, GFHK99wt virus genome replication in the cells was enhanced during coinfection with MaMN99var and NL09var viruses, but not with GFHK99var or GFHK99 PA K26E viruses (Fig 5). However, GFHK99wt virus genome incorporation into progeny virions was enhanced during coinfection with all coinfecting viruses tested. In fact, enhancement relative to the GFHK99wt only control was greater when considering GFHK99wt genomes in progeny virions than when considering their levels within cells (Fig 5). Thus, the beneficial virus-virus interactions observed within cells extend to the release of progeny virions and appear to do so with a higher than expected efficiency.

**Fig 5.**
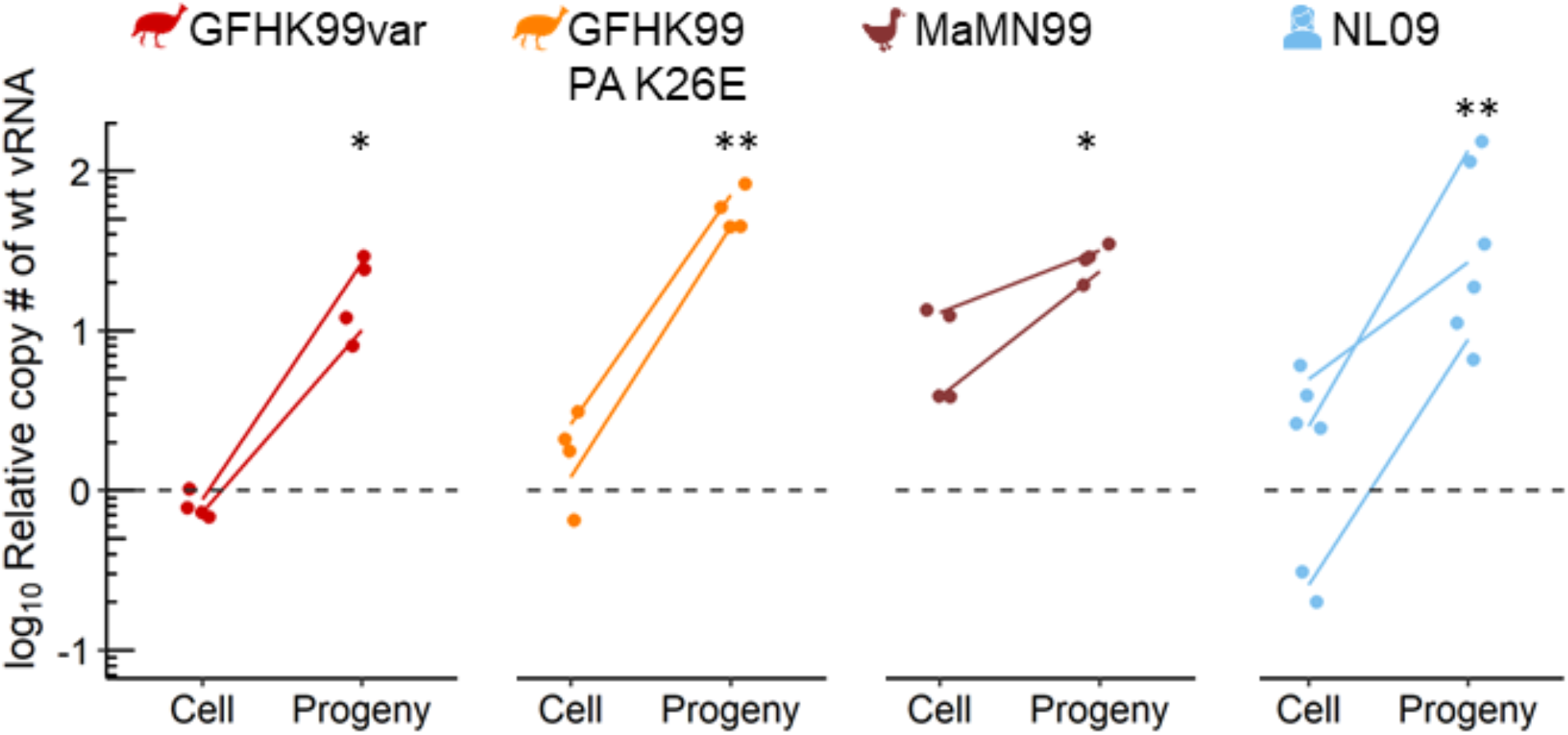
GFHK99 genome incorporation into progeny virions is enhanced during homologous and heterologous coinfection. MDCK cells were coinfected with GFHK99wt virus at an MOI of 1 GC/cell and a homologous GFHK99var or heterologous NL09, MaMN99, or GFHK99var PA K26E virus strains at an MOI of 8 GC/cell. The fold change in wt vRNA copy number, relative to GFHK99wt-only controls (dashed line), in cells and progeny virions collected in the supernatant, is plotted. Results of four to six biological replicates derived from two to three independent experiments are plotted and solid lines connect paired means from each experiment. Significance of differences between wt-only and coinfecting viruses were evaluated by two-way ANOVA and sidak post-hoc analysis; *p< 0.015, **p< 0.01.

### Asynchrony of coinfection and competition for target cells contribute to suppressive virus-virus interactions

The cell culture-based experiments outlined above were all performed under conditions that limited viral replication to a single round, an approach that allows tight control of MOI and detection of intra-cellular virus-virus interactions. Concomitantly, this strategy eliminates the potential for coinfecting viruses to modulate each other’s dynamics during spread through the cellular population. To better understand the dynamics observed in vivo, we therefore adopted a cell culture model in which multiple cycles of viral replication could occur, thereby allowing virus-virus interactions at this higher spatial scale.

Thus, cells were infected at low MOIs in medium conducive to multicycle replication. GFHK99wt virus was used at an MOI of 0.005 GC/cell and GFHK99var or GFHK99 PA K26E virus at an MOI of 0.1 GC/cell. GFHK99wt virus replication was monitored up to 48 h post infection. Consistent with minimal interaction between these relatively small viral populations early in the infection, GFHK99wt virus replication was comparable with or without coinfection up to 24 h post infection (Fig 6 A, B). However, by 36 h or 48 h post infection, a suppressive effect of coinfection was observed (Fig 6 A, B). These data are consistent with observations in guinea pigs, where GFHK99wt replication in coinfected animals begins to show evidence of suppression at 2 days post inoculation (Fig 4A, B). Together, these data suggest that, during multi-cycle replication, limited availability of susceptible target cells gives rise to potent competition between viruses.

**Fig 6.**
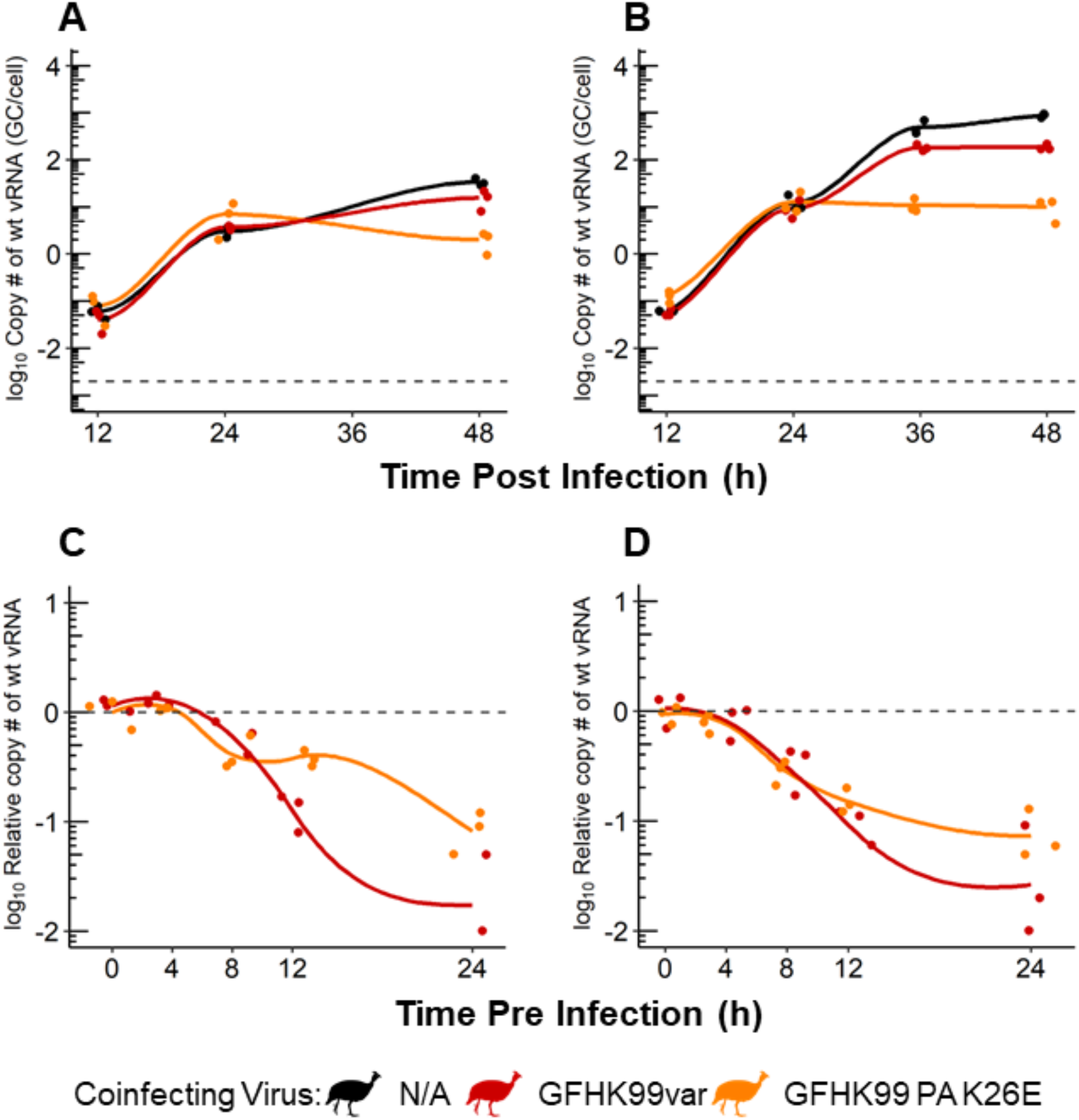
Competitive virus-virus interactions dominate during viral propagation between cells. A-B) MDCK cells were coinfected, under multi-cycle conditions, with GFHK99wt virus at an MOI of 0.005 genome copies GC/cell and GFHK99var or GFHK99 PA K26E virus at an MOI of 0.1 GC/cell. The titer of GFHK99wt virus is plotted and the limit of detection is indicated by the dashed line. C-D) MDCK cells were preinfected with GFHK99var or GFHK99 PA K26E virus, 0–24 h prior to inoculation with GFHK99wt virus. Cells were infected, under single cycle conditions, at an MOI of 0.005 GC/cell of GFHK99wt virus and an MOI of 1 (C) or 10 (D) GC/cell of GFHK99var or GFHK99 PA K26E virus. The fold change in GFHK99wt vRNA copy number, relative to GFHK99wt-only (dashed line), is plotted. Results of three biological replicates derived from one experiment are plotted and solid lines connect the means.

To further evaluate this concept, we tested the impact of modulating the timing of coinfection on the replication of a focal virus. Here, cells were pre-inoculated at varying times up to 24 h before infection with GFHK99var or GFHK99 PA K26E virus at a relatively high MOI and with GFHK99wt virus at low MOI. The replication of the focal virus, GFHK99wt, was then assessed. This experimental design was chosen to model a situation in which most cells were infected (or indirectly affected) by the coinfecting virus prior to introduction of the focal virus. Thus, the potential for cellular coinfection would be high only if cells remain susceptible to infection at the time of GFHK99wt virus introduction. The results show little change to GFHK99wt replication when the coinfecting virus was added up to 4 h before GFHK99wt virus. However, there is a significant decrease in GFHK99wt replication as the interval between infections was increased (Fig 6 C, D). These data indicate that the time window during which beneficial interactions within cells can occur is narrow (between 4 h and 8 h in this system). This temporal effect is likely to be important in shaping viral dynamics during multi-cycle replication, where the introduction of coinfecting viruses into a cell is likely to be asynchronous.

## Discussion

Our results reveal that, within cells, IAV coinfection is strongly beneficial, irrespective of the coinfecting virus’s native host or its homology to the focal strain. Not all coinfecting viruses confer an equal benefit, however, and the magnitude of the benefit is defined by the coinfecting strain’s intrinsic reliance on multiple infection. Despite this clearly cooperative effect within cells, virus-virus interactions at the level of the whole host are antagonistic. Examination of this effect in cell culture suggests that this negative interaction arises at least in part because coinfecting viruses access potential target cells asynchronously, leading to competition for a limited supply of susceptible cells.

Our data suggest a model in which dominant virus-virus interactions differ at the within-cell and within-host scales (Fig 7). Within cells, the positive density dependence of IAV replication extends to heterologous coinfections. However, between cells, the more abundant virus will typically suppress the propagation of the less abundant virus, likely both through depletion of susceptible target cells and triggering of host antiviral responses. This suppression in turn will limit potential for cellular coinfection involving distinct strains. In a natural coinfection, differences in viral abundance are likely to be the norm, resulting from differential fitness, differing initial doses or differing times of introduction into the host. Thus, while intra-cellular interactions within a single viral population are fundamentally important for efficient IAV propagation (14–16, 23), intra-cellular interactions between distinct IAVs are likely to occur more rarely and have a relatively minor effect on within-host viral dynamics. At the level of the whole host, the dominant interaction between distinct viral populations appears to be antagonistic.

**Fig 7.**
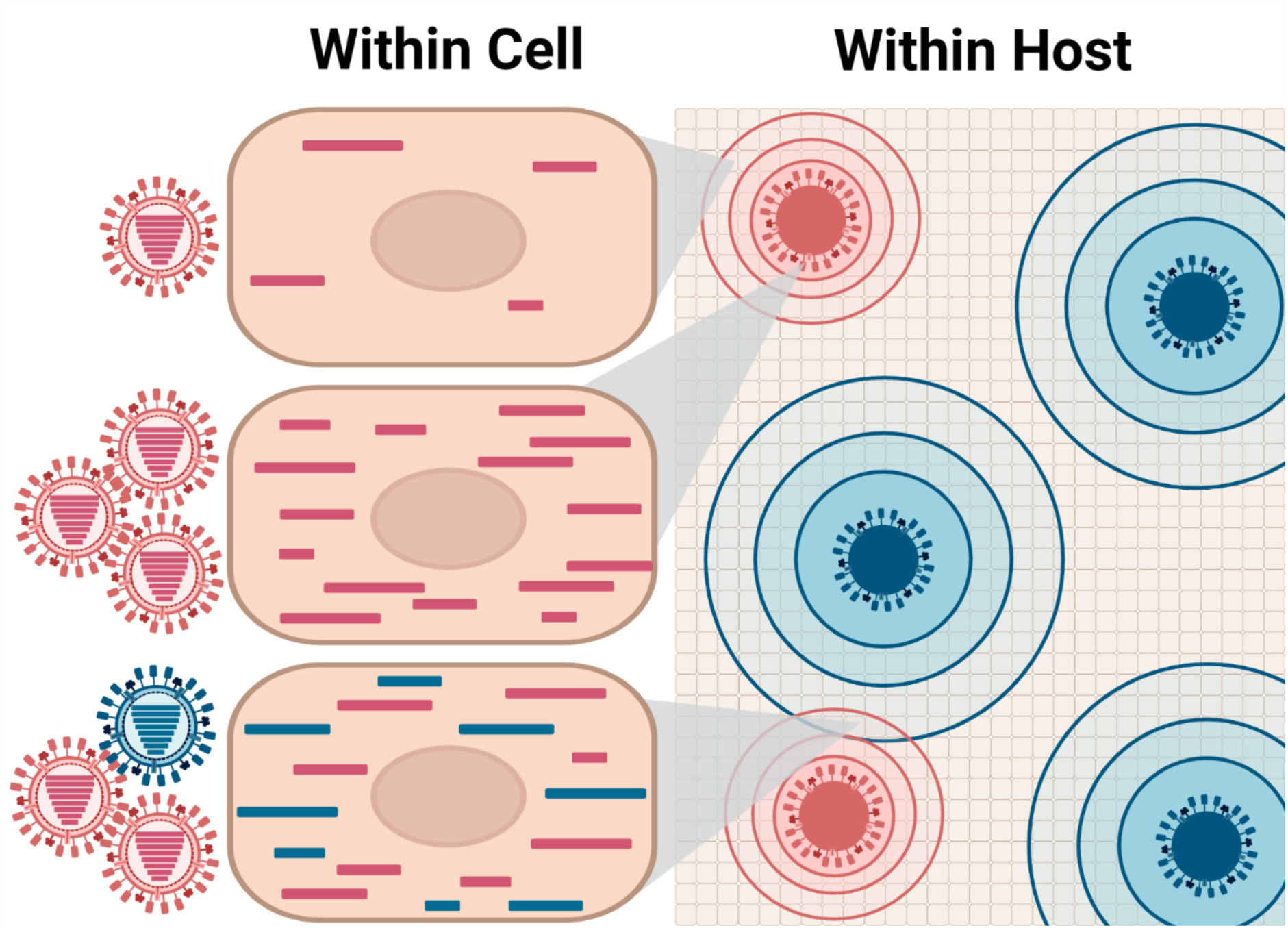
Dominant virus-virus interactions differ at the within-cell and within-host scales. Within an infected cell, coinfection with either homologous (pink) or heterologous (blue) IAVs enhances the replication and augments the progeny production of a focal virus (pink). However, at the within host scale, virus-virus interactions are largely antagonistic as distinct IAV populations compete for susceptible target cells. Target cells can be rendered non-susceptible through super infection exclusion or as a result of cellular antiviral responses triggered indirectly. Within foci, homologous coinfection is highly likely to occur and will promote viral replication. Mixed foci are relatively rare, however, owing to differences in initial dose, the timing of infection or intrinsic fitness leading to discordance in viral population sizes.

For IAVs, cellular coinfection is beneficial in large part because it allows complementation of (very common) incomplete viral genomes, increasing the likelihood of productive infection (14, 15, 24). In addition, cellular coinfection can augment viral yield from productively infected cells by increasing the efficiency of core viral processes (17). Localized viral spread yields areas of high MOI, providing ample opportunity for coinfection, complementation and enhanced replication (15, 21).

At the within-host scale, our data reveal a competitive effect wherein the virus introduced at a higher dose or an earlier time point suppresses the replication of the other. Similar dynamics have been reported previously. In particular, superinfection exclusion, in which the replication of the secondary virus diminished if it is introduced outside of a somewhat narrow time window, is well documented. The duration of the window observed here (4-8 h) is in line with previous work (20, 22, 23, 25). Competitive interactions have also been noted in the context of experiments designed to evaluate the relative fitness of two viral variants (26–30). Conversely, in our own prior work, we have frequently examined coinfection between wt and var viruses with well-matched fitness that are introduced simultaneously and at the same dose. In this scenario, wt and var gene segments are maintained at comparable frequencies throughout the course of infection (19, 22, 31–33). The juxtaposition of these disparate coinfection dynamics suggests that virus-virus interactions within a host are very sensitive to differences between coinfecting strains in their fitness, timing of infection or population size. Effectively, when co-inoculated at equivalent dose, wt and var behave as a single viral population. By contrast, when divergent strains are introduced or when homologous viruses are introduced independently, one strain will typically gain an advantage in the resultant competition for limited resources (or limited time before the immune system responds).

The same conditions that allow the beneficial effects of cellular coinfection to occur also support the replication of defective viral genomes (DVGs). DVGs differ from incomplete viral genomes in that, instead of missing entire segments, DVGs include segment(s) containing large internal deletions. These defective segments can be replicated rapidly and packaged in place of a standard segment, thereby suppressing the production of infectious progeny viruses (34–37). While we ensure low levels of DVGs in our virus stocks and their effects would therefore be minimal in assays limited to a single round of viral replication, we would expect the formation of DVGs *in vivo* (38–40). Prior studies show the presence of DVGs negatively impacts viral replication (13) and can potently activate innate immune responses (41). Thus, delivery of multiple viral genomes to a cell may be predominantly beneficial early on during infection but may become detrimental as DVGs accumulate. In the context of heterologous coinfection, the DVGs of the more abundant coinfecting virus may furthermore drive the suppression of the less abundant strain. While we did not monitor DVG levels in the experiments reported here, these are hypotheses that can be addressed in future work.

Due to the segmented nature of the IAV genome, cellular coinfection yields progeny containing gene segments from both parent strains. Such reassortment plays an important role in IAV evolution in many contexts but is especially prominent in IAV expansion into new host species, including humans (8, 12, 42, 43). The frequency of IAV reassortment involves virus-virus interactions at every biological scale. Exchange of viral gene segments can only occur – and is highly efficient – in coinfected cells (22, 24). Our finding herein that the benefits of cellular coinfection are high even for poorly matched virus strains indicates that virus-virus interactions within cells will promote reassortment between IAVs of distinct lineages. However, our observation that virus-virus interactions at the level of the whole host are antagonistic suggests that the potential for cellular coinfection between independently introduced and/or phylogenetically distinct strains is likely to be limited. When IAVs derived from distinct host species coinfect, the virus replicating in its native host is likely to have a fitness advantage (1, 6, 44, 45). However, if the host has pre-existing immunity to the well-adapted strain, this could tip the balance in favor of the novel virus. Of course, should conditions of timing, dose and fitness combine to allow reassortment, a chimeric strain that brings together genes well-adapted to the host with genes encoding novel antigenic determinants can result. In humans, such a virus would have pandemic potential. Our data nonetheless suggest that the propagation of such a strain within the host would usually be strongly limited by the already established parental virus populations. Onward transmission, in turn, would be unlikely owing to tight transmission bottlenecks (46–48). In sum, the antagonistic nature of virus-virus interactions at the within host scale is likely a major factor contributing to the rarity of IAV pandemics.

In conclusion, while the benefits of cellular coinfection extend to heterologous strains, disparate fitness or conditions of introduction into a host will typically limit the opportunity for heterologous strains to meet within cells. The need for multiple infection is therefore more likely to be met within a single viral population. Nevertheless, IAV populations coinfecting the same host have a significant impact on each other’s dynamics owing to competition for limited resources.

## Author Contributions

Concept and experimental planning was performed by GD and ACL. Data was collected GD, KG and KEH and analyzed by GD with input from ACL. Key reagents and intellectual input were provided by JRS. Manuscript and figures were written, designed, and edited by GD and ACL. Research funding was acquired by ACL.

## Acknowledgements

This work was supported by funding from NIH/NIAID under the Centers of Excellence for Influenza Research and Response contract no. 75N93021C00017 to ACL and R01 AI127799 to ACL.

## Materials and Methods

### Ethics statement

Experiments using guinea pigs were conducted in accordance with the Guide for the Care and Use of Laboratory Animals of the National Institutes of Health. All studies were approved by the Emory University Institutional Animal Use and Care Committee (protocol PROTO201700595) and were conducted under animal biosafety level (ABSL-2) containment. Guinea pigs were humanely euthanized following America Veterinary Medical Association approved guidelines.

### Cells

Madin-Darby Canine Kidney (MDCK) cells, a gift of Daniel Perez at University of Georgia, were maintained in minimum essential medium (MEM; Gibco) supplemented with Normocin (InvivoGen) and 10% Fetal Bovine Serum (FBS). Human kidney 293T cells (ATCC CRL-3216) were maintained in Dulbecco’s minimal essential medium (Gibco) supplemented with Normocin and 10% FBS. Normal Human Bronchial Epithelial (NHBE) cells (Lonza) were maintained in bronchial epithelial cell growth medium (BEGM) purchased from Lonza. NHBE cells from a single donor were amplified and differentiated into air-liquid interface cultures as recommended by Lonza and described in (49). All cells were cultured at 37°C and 5% CO_2_ in a humidified incubator. Cells were tested monthly for mycoplasma contamination during use. Medium used for IAV infection in each cell line (virus medium) was prepared using the appropriate medium containing Normocin and 4.3% bovine serum albumin. Infection of NHBE cells was performed with BEGM in the basolateral chamber. Virus medium or BEGM containing ammonium chloride was prepared by adding HEPES buffer and NH4Cl at final concentrations of 50 mM and 20 mM, respectively.

### Viruses

The strains influenza A/Netherlands/602/2009 (H1N1) and A/Panama/2007/99 (H3N2) are referred to herein as NL09 and Pan99, respectively. NL09 and Pan99 were handled under BSL2 conditions. The strains influenza A/guinea fowl/Hong Kong/GFHK99/99 (H9N2) and A/mallard/Minnesota/199106/99 (H3N8) are referred to herein as GFHK99 and MaMN99, respectively. GFHK99 and MaMN99 were handled under BSL2 conditions with enhancements as required by the United States Department of Agriculture. Coinfections involving human and avian IAVs were performed under BSL3 conditions.

All viruses used were generated using reverse genetics (50, 51). The reverse genetics system for GFHK99 is in the ambisense vector pDP and was a kind gift of Daniel Perez. The reverse genetics system for NL09 is in the ambisense vector pHW and was a kind gift of Ron Fouchier. The reverse genetics systems for Pan99 and MaMN99 viruses were generated in house in the pDP vector (a gift of Daniel Perez). The GFHK99 PA K26E mutant virus was described in (17) and was generated by introducing the K26E mutation to the reverse genetics plasmid encoding the PA gene segment of GFHK99 using site-directed mutagenesis. The specific mutations introduced are listed in S1 Table. Silent mutations were introduced into var viruses by site-directed mutagenesis to allow discrimination between wt and var gene segments by site-specific primers used in ddPCR. The specific mutations introduced into all var viruses are listed in S1 Table.

Briefly, avian viruses were generated by transfecting 293T cells with eight reverse genetics plasmids encoding each IAV segment. After 16 h, transfected 293T cells were injected into the allantoic cavity of 11-day old chicken eggs, incubated at 37°C for 40-48 h, and allantoic fluid was recovered for use as a passage 1 working virus stock. Mammalian IAVs were generated by transfecting 293T cells with reverse genetics plasmids encoding each IAV segment. After 16 h, transfected 293T cells were then cocultured with MDCK cells at 37°C for 40-48 h. Collected supernatants were then propagated in MDCK cells from low MOI to generate a working virus stock. Every virus stock was tested for defective viral genomes (DVGs) as described previously (52) (S4 Fig). We define ‘low DVG content’ as follows: the ratio of copy number for terminal and internal targets (T:I) is <2.0 for each of the PB2, PB1 and PA gene segments. Any stocks not meeting this criterion were regenerated.

Viral concentrations are reported in genome copies per mL throughout this work for two reasons. First, use of a molecular assay for viral quantification allows differentiation of IAVs present in a mixture. Second, this approach ensures consistency in the limit of detection across virus strains. In this work and in prior studies, we have found that viral infectivity varies with MOI and the efficiency with which a single virus particle initiates infection varies widely across strains. As a result, titration of infectious units at limiting dilution by plaque assay or TCID_50_ assay measures differing sub-populations for different IAV strains.

### Synchronized, single- and multi-cycle infections

Synchronization was used to coordinate the timing of viral entry into cells and was carried out as follows: The monolayer of cells was washed 3x with cold PBS and placed on ice. Chilled virus inoculum was added to each well and kept at 4°C for 45 min with rocking to allow time for IAV attachment but not entry. After that time, the inoculum was aspirated, each well was washed 3x with cold PBS. Warmed virus medium without trypsin was added to allow entry and this time is defined as t=0. Cultures were incubated at 37°C. Single-cycle conditions were designed to prevent released progeny virus from initiating a subsequent round of infection. A single cycle of infection was imposed by replacing virus medium with virus medium containing NH4Cl and HEPES solution at final concentrations of 20 mM and 50 mM, respectively, at 3 h post-infection. In the case of NHBE cells, this medium was added to the basolateral chamber only. Addition of NH4Cl and HEPES to the medium prevents the acidification of endosomes, thereby blocking infection (42). For multi-cycle infections, L-1-tosylamido-2-phenylethyl chloromethyl ketone (TPCK)-treated trypsin was added to virus medium at 0 h post-infection at a final concentration of 1 μM, and NH4Cl and HEPEs were not added. Addition of TPCK-treated trypsin allows cleavage activation of the HA protein on released virus, which is required for infection.

### Detection of virus-virus interactions within cells

MDCK cells were seeded into 12-well plates at 4 × 10^5^ cells per well 24 h prior to infection. NHBE cells were cultured at an air-liquid interface as previously described (49). Each infection took place under synchronized, single-cycle conditions. For homologous coinfections, triplicate wells were inoculated with wt virus at an MOI of 0.005 genome copies (GC)/cell in MDCK cells, or 0.5 GC/cell in NHBE cells and increasing doses of a matched var virus. For heterologous coinfections, triplicate wells were inoculated with GFHK99wt virus at an MOI of 0.005 GC/cell in MDCK cells, or 0.5 GC/cell in NHBE cells, and increasing doses of MaMN99wt, NL09var, or Pan99wt viruses. At 12 h post infection, virus medium containing NH4Cl and HEPES was removed, cell surfaces were washed 3x with PBS, and cells were harvested for RNA extraction using the Qiagen RNeasy mini kit.

### Impact of intracellular virus-virus interactions on released viral progeny

MDCK cells were seeded into T75 flasks at 5 × 10^6^ cells per flask 24 h prior to infection. Each infection took place under synchronized, single-cycle conditions. Duplicate flasks were inoculated with GFHK99wt virus at an MOI of 1 GC/cell and either PBS, homologous GFHK99var virus, or heterologous MaMN99wt, NL09var, or Pan99wt viruses at an MOI of 8 GC/cell. At 24 h post infection medium (containing viral progeny) was collected separately from cells. Cells were washed 3x with PBS and harvested using the Qiagen RNeasy mini kit and protocol instructions. Collected medium was centrifuged (Thermofisher Sorvall ST 16R Centrifuge) at 3000 rpm for 10 min at 4°C to remove cell debris. The resultant supernatant was transferred into a Beckman unltracentrifuge tube and centrifuged (Beckman Coulter Optima™ XL-100K Ultracentrifuge) at 10,000 rpm for 30 min at 4°C to further clarify the sample of cell debris. The final supernatant was then transferred into a new unltracentrifuge tube. A cushion of 5 mL 30% sucrose in NTE buffer[1 M NaCl, 0.1 M Tris, 0.01 M EDTA, pH 7.4] as injected into the bottom of the tube and the sample was centrifuged (Beckman Coulter Optima™ XL-100K Ultracentrifuge) at 25,000 rpm for 2 h at 4°C. The resultant viral pellet was resuspended in PBS and RNA extracted using the Qiagen Viral RNA mini kit and protocol instructions.

### Monitoring virus-virus interactions in guinea pigs

Female Hartley strain guinea pigs weighing 300-350 g were obtained from Charles River Laboratories (Wilmington, MA) and housed by Emory University’s Department of Animal resources. Prior to intranasal inoculation, nasal lavage and euthasia, guinea pigs were anesthetized by intramuscular injection with 30 mg/kg ketamine and 4 mg/kg xylazine. Guinea pigs were inoculated intranasally with IAV in 300 μl PBS. Daily nasal washes were performed up to 6 d post GFHK99wt inoculation. Briefly, with the animal’s nose suspended in a downward orientation over a Petri dish, 1 mL PBS was instilled into the nares in 200 μl increments and allowed to drop back out. Liquid was pooled into a 1.5 mL tube, aliquoted and stored at - 80°C.

### Quantification of vRNA

RNA was extracted from cells and viral samples using the Qiagen RNeasy mini kit and the Qiagen Viral RNA mini kit, respectively, using included protocol instructions. Extracted vRNA was reverse transcribed using a 1:1 ratio of universal influenza primers (53)(S2 Table) and Maxima RT (Thermofisher) per protocol instructions. Droplet digital PCR (ddPCR) was performed on the resultant cDNA using the QX200™ ddPCR™ EvaGreen Supermix (Bio-Rad) and virus specific primers targeting the NP segment (final concentration 200 nM) (S2 Table). Copy numbers of the wt virus present in coinfected samples were normalized to those detected in wt only control infections.

### Statistical analyses

All statistical analyses were performed in R (version 1.3.959). Log transformed values for genome copy numbers and the mean of technical replicates were used for statistical analyses.

