## Supplemental Materials for "Influenza A virus coinfection dynamics are shaped by distinct virus-virus interactions within and between cells"

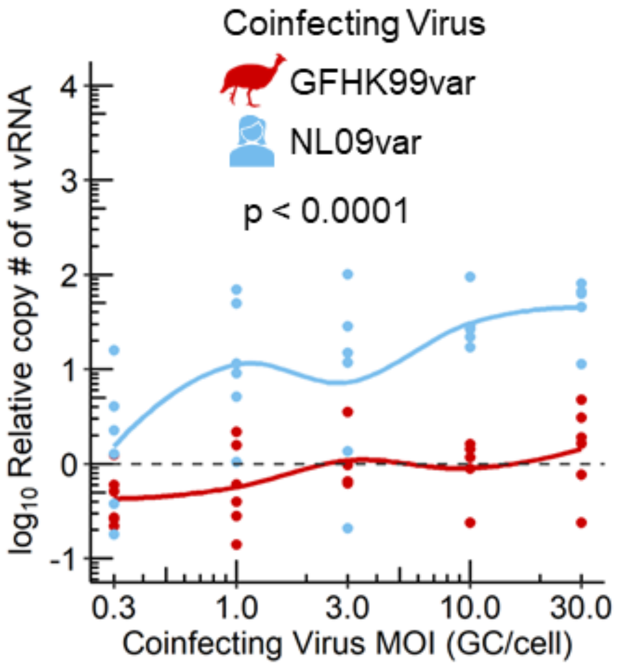


**S1 Fig. Coinfection with NL09 strongly enhances GFHK99wt replication in NHBE cells.** (Related to Fig 2.) NHBE cells were infected with GFHK99wt virus at an MOI of 0.5 genome copies (GC)/cell and increasing doses of the homologous GFHK99var virus or the heterologous NL09 virus. The fold change in GFHK99wt vRNA copy number, relative to GFHK99wt-only control (dashed line), is plotted. Results of six biological replicates derived from two independent experiments are plotted and solid lines connect the means. Significance of differences between results obtained with the differing coinfecting viruses were evaluated by two-way ANOVA.

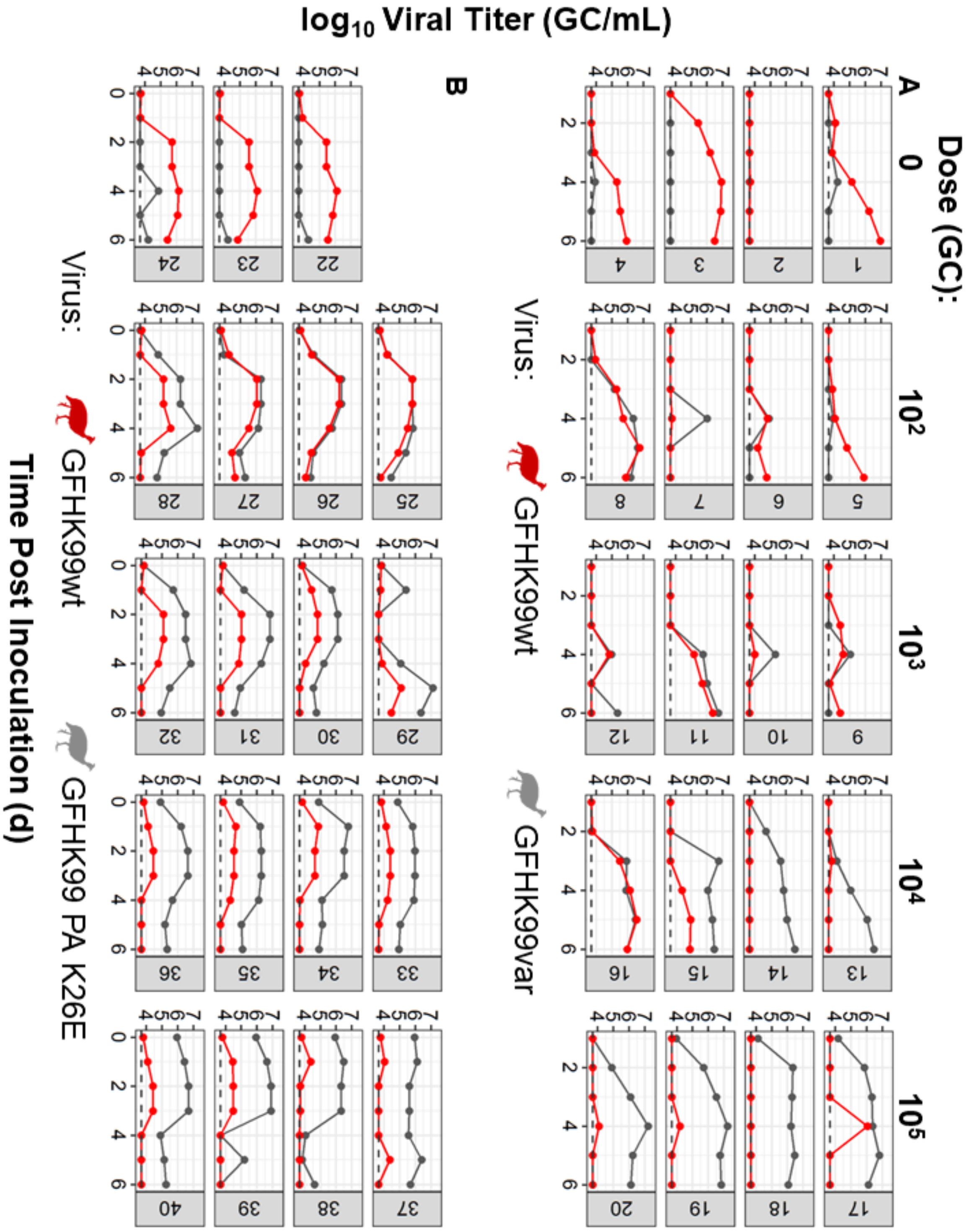


**S2 Fig. At the level of the whole host, GFHK99 replication is suppressed by coinfection.** (Related to Fig 5A). Guinea pigs were infected with GFHK99wt at a dose of $10^3$ GC and increasing doses of GFHK99var (A) or GFHK99 PA K26E (B) virus. The viral titer in nasal washes collected from each guinea pig is plotted and the limit of is indicated by the dashed line. Guinea pig ID numbers are shown in grey boxes appended to each facet.

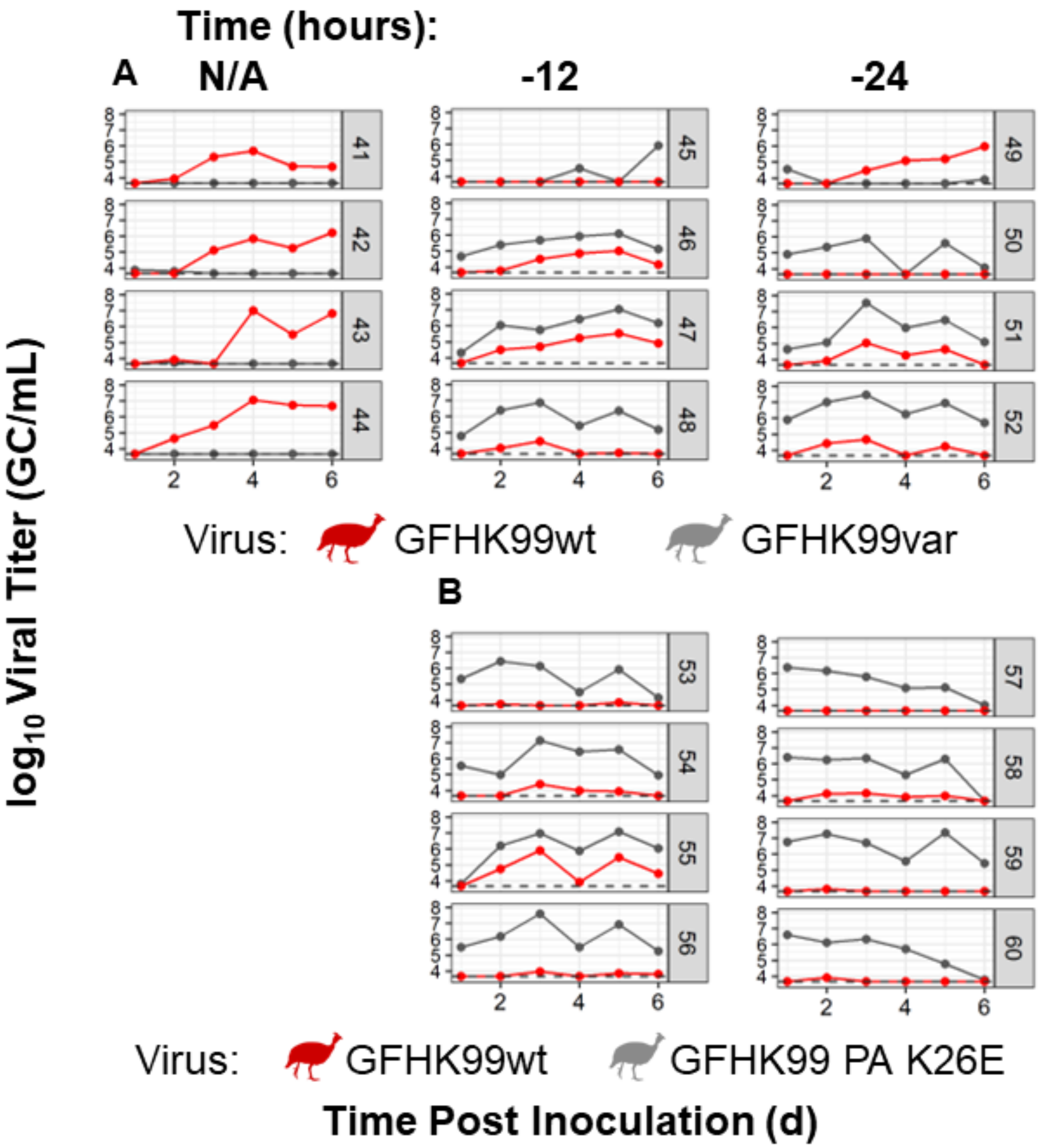


**S3 Fig. At the level of the whole host, GFHK99 replication is suppressed by prior IAV infection.** (Related to Figure 4B). Guinea pigs were pre-inoculated with $10^4$ GC of either GFHK99var (A) or GFHK99 PA K26E (B) virus either 12 or 24 h prior to a dose of $10^4$ GC of GFHK99wt virus. The viral titer in nasal washes is plotted and the limit of detection is indicated by the dashed line. Guinea pig ID numbers are shown in grey boxes appended to each facet.

**S1 Table. Genotypes of modified viruses.**

| | PB2 | PB1 | PA | HA | NP | NA | M | NS |
|---|---|---|---|---|---|---|---|---|
| **GFHK99var$_2$** | A300G, A303T, T306C, G459C, T461A, T467T | T282C, T285C, A288G, A420G, T426C, C432T | A351G, C354T, C357T, T501G, C504T, C507T | A338G, A344C, A351C, A432G, T435A, C438T | A345G, G351A, A354G, A486T, C489T, A495G | A424G, T430A, G433A, A583G, G586C, G589C | G340A, A343G, A349G, G439A, C442T, A445G | C386T, G389A, A392G, A479G, G482C, A488G |
| **NL09var** | C273T | T288C | C360T | C305T | A351G | G336A | G295A | C341T |
| **Pan99var0** | A345T, C360T | A540G | G333A, A342G | T308A, C311A, C314T, A464T, C467G, T470A | C537T, T538A, C539G, C612G, G615A | C418G, T421A, A424C | G586A | A329T, A335T, C341G |
| **GFHK99wt PA K26E** | --- | --- | A100G, A102G | --- | --- | --- | --- | --- |
| **GFHK99var$_2$ PA K26E** | A300G, A303T, T306C, G459C, T461A, T467T | T282C, T285C, A288G, A420G, T426C, C432T | A100G, A102G, A351G, C354T, C357T, T501G, C504T, C507T | --- | A345G, G351A, A354G, A486T, C489T, A495G | A424G, T430A, G433A, A583G, G586C, G589C | G340A, A343G, A349G, G439A, C442T, A445G | C386T, G389A, A392G, A479G, G482C, A488G |

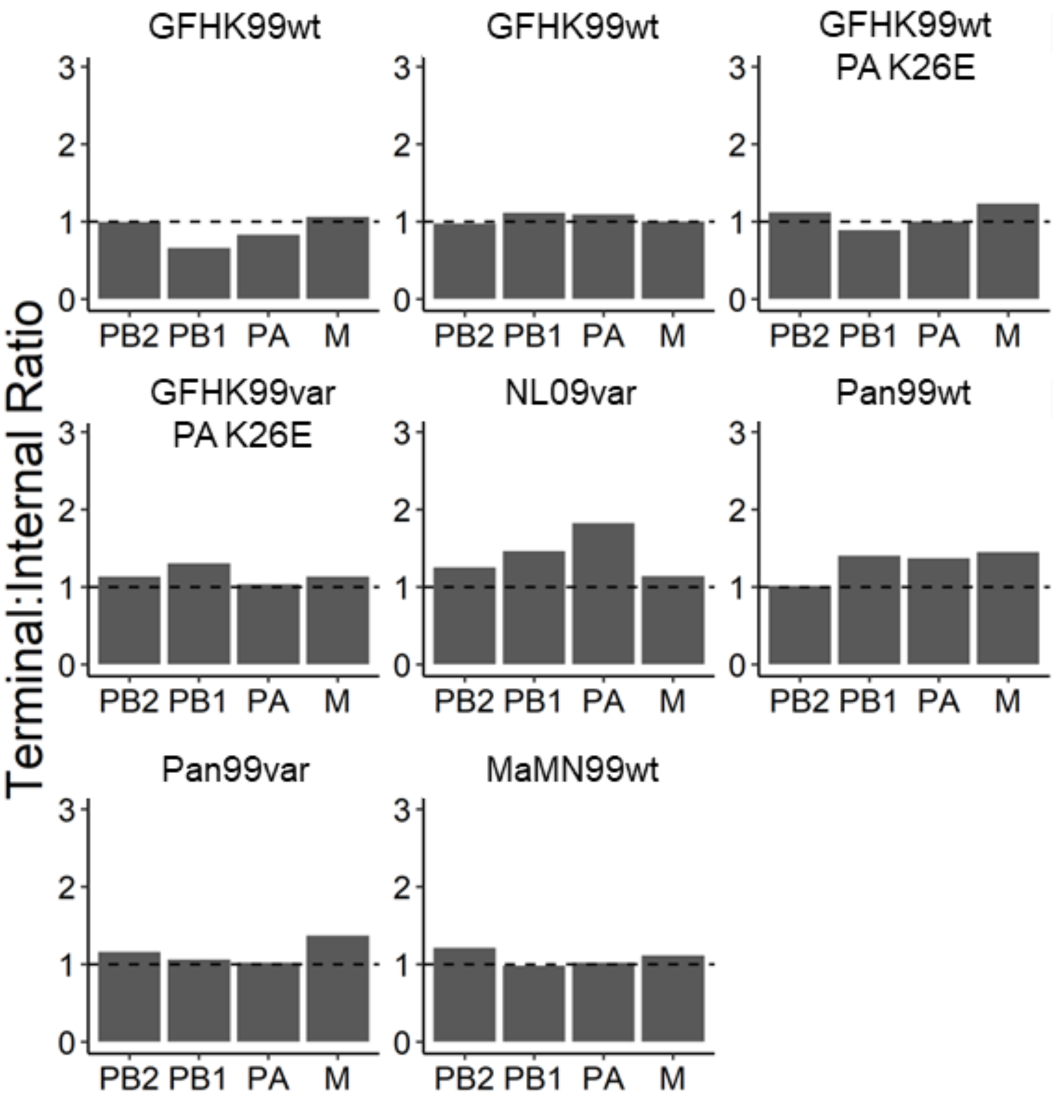


**S4 Fig. Quantification of defective viral genomes in virus stocks.** Levels of defective viral genomes (DVGs) were quantified by ddPCR using primers targeting the terminal and internal regions of the PB2, PB1, PA, and M gene segments. Ratios of terminal to internal copies that are <2.0 indicate low DVG content. Virus stocks tested are named above each facet. Two different stocks of GFHK99wt virus were used.

**S2 Table. Primers for the quantification of vRNA by ddPCR.**

| | |
|---|---|
| **Universal influenza Reverse Transcription Primers** | |
| Univ.F(A)+6 | GCGCGCAGCAAAAGCAGG |
| Univ.F(G)+6 | GCGCGCAGCGAAAGCAGG |
| **GFHK99wt, GFHK99wt PA K26E Virus Primers** | |
| WF10wt NP 336 F | GAAGGAGAGACGGGAAATG |
| WF10wt NP 505 R | GGCTCTTGTTCTCTGGTATG |
| **GFHK99var$_2$, GFHK99var$_2$ PA K26E Virus Primers** | |
| WF10help NP 388 F | GAAGGAGGGACGGAAAGT |
| WF10help NP 505R | GGGCTCTTGTCCTCTGATAA |
| **NL09var Virus Primers** | |
| NL09 NP 309 F | CCCTAAGAAAACAGGAGGACCC |
| NL09 NP 411 R | TTGGCGCCAAACTCTCCTTA |
| **Pan99wt Virus Primers** | |
| Pan99wt NP 520 F | ATGGATCCCAGAATGTGCTC |
| Pan99wt NP 625 R | TCAGCTCCATTGTC |
| **Pan99var Virus Primers** | |
| Pan99var0 NP 520 F | ATGGATCCCAGAATGTGTAG |
| Pan99var0 NP 625 R | TCAGCTCCATAGTG |
| **MaMN99wt Virus Primers** | |
| MN99 NP 378 F | CGACAAAGAAGAGATCAGAAGGA |
| MN99 NP 457 R | TCATCAAATGGGTGAGACCA |
